# Segregational drift and the interplay between plasmid copy number and evolvability

**DOI:** 10.1101/369579

**Authors:** Judith Ilhan, Anne Kupczok, Christian Woehle, Tanita Wein, Nils F. Hülter, Philip Rosenstiel, Giddy Landan, Itzhak Mizrahi, Tal Dagan

## Abstract

The ubiquity of plasmids in all prokaryotic phyla and habitats and their ability to transfer between cells marks them as prominent constituents of prokaryotic genomes. Many plasmids are found in their host cell in multiple copies. This leads to an increased mutational supply of plasmid-encoded genes and genetically heterogeneous plasmid genomes. Nonetheless, the segregation of plasmid copies into daughter cells during cell division is considered to occur in the absence of selection on the plasmid alleles. We investigate the implications of random genetic drift of multicopy plasmids during cell division – termed here *segregational drift* – to plasmid evolution. Performing experimental evolution of low- and high-copy non-mobile plasmids in *Escherichia coli*, we find that the evolutionary rate of multicopy plasmids does not reflect the increased mutational supply expected according to their copy number. In addition, simulated evolution of multicopy plasmid alleles demonstrates that segregational drift leads to increased loss frequency and extended fixation time of plasmid mutations in comparison to haploid chromosomes. Furthermore, an examination of the experimentally evolved hosts reveals a significant impact of the plasmid type on the host chromosome evolution. Our study demonstrates that segregational drift of multicopy plasmids interferes with the retention and fixation of novel plasmid variants. Depending on the selection pressure on newly emerging variants, plasmid genomes may evolve slower than haploid chromosomes, regardless of their higher mutational supply. We suggest that plasmid copy number is an important determinant of plasmid evolvability due to the manifestation of segregational drift.

## Introduction

Plasmids are genetic elements that colonize prokaryotic cells where they replicate independently of the chromosome. They are considered as a major driving force in prokaryotic ecology and evolution as they can be transferred between cells, making them potent agents of lateral gene transfer (Lederberg and Tatum 1946; Thomas and Nielsen 2005) and microbial warfare (Czárán et al. 2002; Majeed et al. 2011). Plasmids may contribute to the evolution of their host via the supply of genes encoding for new functions that are beneficial under growth limiting conditions (e.g., antibiotics resistance (Porse et al. 2016), resistance to heavy metals (Gullberg et al. 2014; Dziewit et al. 2015)) or during adaptation to new habitats (e.g., catabolic functions (Schmidt et al. 2011)). Nonetheless, the importance of plasmids goes beyond microbial evolution as they are widely used as vectors for genetic engineering (Simon et al. 1983; Bevan 1984) as well as for applications in biotechnology (Ullrich et al. 1977) and synthetic biology (Shetty et al. 2008). Therefore, understanding plasmid biology and evolution is of major interest for basic microbiology research and biological applications.

Plasmids differ from the chromosomal genetic component in several key aspects, including a relatively small genome size and a variable ploidy level. Plasmids are often found in multiple copies in the cell where the plasmid copy number (PCN) depends on the replicon type, the host genetics, and the environmental conditions (Nordström 2006; Santos-Lopez et al. 2017). The copy number of natural plasmids ranges between one and 15 for low-copy plasmids (Bazaeal and Helinski 1968) and can reach 200 copies for high-copy plasmids (Projan et al. 1987). Plasmids used for biotechnological applications are often modified to have a higher copy number, e.g., 500 to 700 (Vieira and Messing 1982; Rosano and Ceccarelli 2014), in order to maximize the protein expression level of focal genes. Notably, genes encoded on multicopy plasmids residing within a single host cell may comprise multiple alleles due to independent emergence of mutations in the plasmid copies (Bedhomme et al. 2017; Rodriguez-Beltran et al. 2018). An increased plasmid copy number has the potential to elevate the probability for the emergence of novel mutations in the gene open reading frame. Consequently, the mutational supply of plasmid-encoded genes is increased for multicopy plasmids. Indeed, it has been previously shown that multicopy plasmids have the potential to enable rapid evolution of plasmid-encoded antibiotic resistance genes under strong selective conditions for the antibiotic resistance (San Millan et al. 2016).

The inheritance of multicopy plasmids to daughter cells depends on their segregation mechanism. Some plasmids are equipped with an active partition system. Such systems comprise plasmid-encoded proteins that function in the translocation of plasmid copies toward the cell poles prior to cell division, thereby ensuring stable inheritance (e.g., *parABS* system (Abeles et al. 1985; Baxter and Funnell 2014)). In the absence of a partition system, plasmid segregation into daughter cells depends on the physical distribution of plasmid copies in the cell during cell division (Wang 2017). Notably, plasmid segregation into daughter cells in both routes is considered to occur in the absence of selection on the plasmid allele pool. Consequently, the dynamics of multicopy plasmid alleles in the population has a constant component of random genetic drift. In other words, the distribution of plasmid alleles into daughter cells is independent of the allele impact on the host fitness, also under selective conditions for a plasmid-encoded trait. Here we term this phenomenon *segregational drift*. Previous studies suggested that segregational drift plays a role in the evolution of eukaryotic organelles – mitochondria and plastids (Birky 2001). Similar to multicopy plasmids, the mitochondrion organelle is found in multiple copies in the cell (e.g., ~1,700 in mammalian cells (Robin and Wong 1988)), and in addition, each mitochondrion organelle may harbour multiple copies of the mitochondrial genome (e.g., up to 10 copies (Satoh and Kuroiwa 1991)). Thus, the mutational supply on the mitochondrial genome is larger than the mitochondrial genome size by several orders of magnitude. The combination of high mutational supply on the mitochondrion genome and random segregation of the mitochondria during cell division lead to genetic heterogeneity of the mitochondrial genomes within a single cell (termed ‘mitochondrion heteroplasmy’ (Birky 2001)). Notably, many of the mitochondrial alleles are found in very low frequency, thus the mitochondrial genetic heterogeneity is observed only with the application of deep-sequencing approaches (e.g., (He et al. 2010)). The consequences of segregational drift to the evolution of multicopy plasmids have been so far overlooked.

Population genetics theory postulates that the effect of random genetic drift on allele dynamics is inversely correlated with the effective population size. In a larger population, the impact of genetic drift on the dynamics of allele frequency is reduced (see (Lanfear et al. 2014) for review). Thus, the implications of segregational drift to plasmid genome evolution are expected to be dependent on the plasmid population size within the cell, i.e., the plasmid copy number. Here we investigate plasmid genome evolution in light of the segregational drift using experimental evolution and *in silico* simulation of allele dynamics in the population.

## Results

### Experimental evolution

In our study we compare the rate of evolution between low-copy and high-copy plasmid replicons using an experimental evolution approach. In the experiment, two model plasmids having low- or high-copy number were evolved in an *E. coli* host under conditions selecting for the plasmid presence. The low-copy model plasmid (pLC, 5.2 kb; Fig. 1) is derived from plasmid pBBR1 (Antoine and Locht 1992) that is known to replicate in a broad host range in the absence of an active partitioning mechanism. The plasmid encodes an ampicillin-resistance gene (*bla*; β-lactamase) and is non-mobile (i.e., it is not transmissible via conjugation). The average pLC copy number in the wild-type *E. coli* host was 8.0 ± 0.78 (*SD*, *n* = 12; using qPCR), which is within the range calculated previously for pBBR1 (Jahn et al. 2016). The accessory part of the high copy number model plasmid (pHC, 4.8 kb; Fig. 1) is identical to that of plasmid pLC, but the backbone comprises a pUC origin of replication (Vieira and Messing 1982), that is a derivative of the *ori* of plasmid ColE1 (Hershfield et al. 1974). The copy number of plasmid pHC in the wild-type *E. coli* host was approximately 80-fold higher than that of plasmid pLC (715.56 ± 27.4 *SD*, *n* = 12; using qPCR). This is within the typical copy number range of the pUC plasmid backbone (Rosano and Ceccarelli 2014). In addition to PCN, our experimental evolution setup accounts for two known determinants of evolutionary dynamics: mutation rate and environmental conditions. To vary the mutation rate, the plasmids were evolved in two host genotypes, *E. coli* K-12 substrain MG1655 (wild-type, wt) and a hypermutator derivative of the wild-type strain (Δ*mutS*) in which mutations are expected to arise faster in comparison to the wild-type (e.g., 33-fold (Arjan et al. 1999)). The populations were evolved under two different environmental regimes: at 37°C and at 42°C. Growth of *E. coli* at 42°C is expected to entail an acclimation to the elevated temperature (Guyot et al. 2014), and it has been shown to involve genomic adaptation (Deatherage et al. 2017) The experiment was conducted with six replicates for each combination of the three experimental evolution factors – plasmid replicon type, host genotype and growth temperature. The populations were evolved for 800 generations in continuous culture under selective conditions for ampicillin-resistance. At the end of the experiment, samples of ancestral and evolved populations were sequenced using a population sequencing approach. Notably, the two model plasmids differ in their stability within the host. While the pLC replicon remained unaltered compared to its pBBR1 ancestor, the high copy number of plasmid pHC comes at the cost of lower plasmid stability due to deletions of stability factors during construction of the pUC replicon (e.g., lack of the multimer resolution site *cer* originally present in plasmid ColE1). We assessed the stability of both plasmids in the *E. coli* wild-type host using the same selective constraints for the plasmid presence as in the evolution experiment. No plasmid-free segregants of pLC-wt were observed after overnight growth (*n* = 5), whereas the proportion of pHC-free segregants ranged between 1% and 39% (*n* = 5; see Methods). Plasmid loss during the evolution experiment has the potential to result in a proportion of plasmid-free cells in the total population. Such plasmid-free cells have a collective resistance to ampicillin thanks to the presence of β-lactamase secreted from plasmid carrying hosts in the environment (Vega and Gore 2014). Therefore, we quantified the proportion of plasmid-hosts in the evolved populations using replica plating of single colonies. This showed that pLC-hosts accounted for 89% to 100% of the populations in all conditions and replicates (Supplementary Table S1). The proportion of pHC-hosts in the evolved wild-type populations ranged between 1% and 78%, while the range of plasmid carrying hosts in the evolved hypermutator populations was between 35% and 98% (Supplementary Table S1). In order to increase the resolution of the existing plasmid allele pool, we sequenced the genomes of each pHC population using two different sampling strategies. In addition to the sequencing of total populations (i.e., plasmid host cells and plasmid-free segregants), the genome of each pHC population was sequenced from pooled samples of 100 colonies of plasmid carrying hosts (see Methods). In the following, the resulting genetic variants reported for pHC populations are based on the pooled colonies data unless stated otherwise.

**Figure 1.**
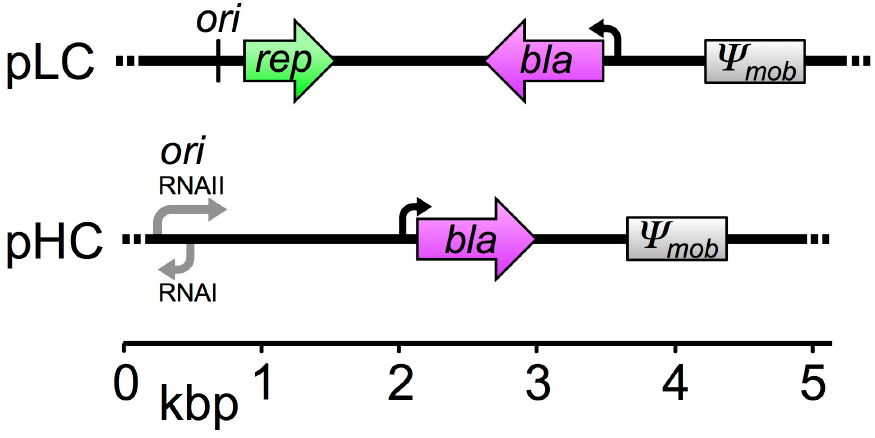
Genetic maps of plasmid pLC and pHC. The plasmids are of comparable genome size and differ in the origin of replication. The *mob* gene originally found in pBBR1 was non-functionalized by truncation during plasmid construction (denoted as *Ψ*_mob_).

### Plasmid genome evolution

For the identification of emerging mutations, sequencing reads of the ancestral and evolved populations were mapped to the reference *E. coli* MG1655 genome and to the two plasmids. The plasmid copy number within the sequenced plasmid host population was estimated from the average sequencing coverage of the plasmids relative to that of the chromosome. The sequencing coverage varied among the populations and the replicons: the chromosomal coverage ranged between 104x and 300x, while the average coverage on the plasmids was between 325x and 28,054x (Supplementary Table S1). The high coverage on the plasmid genomes increases the detection sensitivity for rare and low-frequency variants. Nonetheless, in order to compare the number of mutations across replicons and populations, the sequencing reads of each replicon (i.e., chromosome and plasmids) were subsampled to match the lowest-coverage dataset unless stated otherwise. Furthermore, variants occurring in both ancestral and evolved populations were excluded.

A comparative genomic analysis of the subsampled sequencing data revealed two point mutations on the pLC plasmid (Table 1). Both mutations are located in an intergenic region and their allele frequency (AF) is lower than 0.10 (Supplementary Table S2). The paucity of plasmid variants should be evaluated against the background of variants on the host chromosome, where we observed a total of 1,394 point mutations. We calculated the expected frequency of point mutations on the two plasmids using the chromosomal substitution frequency, the plasmid genome size and the plasmid copy number. This shows that mutations are unlikely to occur on either of the plasmid types evolved in the wild-type host due to the combination of low substitution rate and the plasmid small genome size (Table 1). In contrast, the combination of high genetic polymorphism in the hypermutator host with the high copy number of pHC are likely to result in observable genetic variants in the pHC plasmid within the time span of our experiment (Table 1 and Supplementary Table S1). Nonetheless, the subsampled sequencing data for pHC did not reveal any point mutations occurring on that plasmid. Notably, we detected two single nucleotide insertions in pHC intergenic regions with an allele frequency of AF = 0.3 and AF = 0.35. In order to increase the resolution for the detection of presumably low-frequency point mutations, we applied the same variant calling pipeline to the sequencing read datasets without subsampling (i.e., full sequencing coverage). This analysis revealed one point mutation on plasmid pHC at an intergenic position with a sequencing depth of ~9,600 and a low allele frequency of AF = 0.03 (Supplementary Table S2). In contrast, considering the full sequencing depth for plasmid pLC, which is on average ~10x lower than that of plasmid pHC, we observed two additional intergenic point mutations (Supplementary Table S2). Thus, despite the high sequencing depth, the observed number of point mutations on plasmid pHC does not reach the expected frequency of point mutations, even when the full sequencing coverage is included in the analysis. On the other hand, the number of point mutations detected on plasmid pLC is higher than what we expected (Table 1).

**Table 1.**
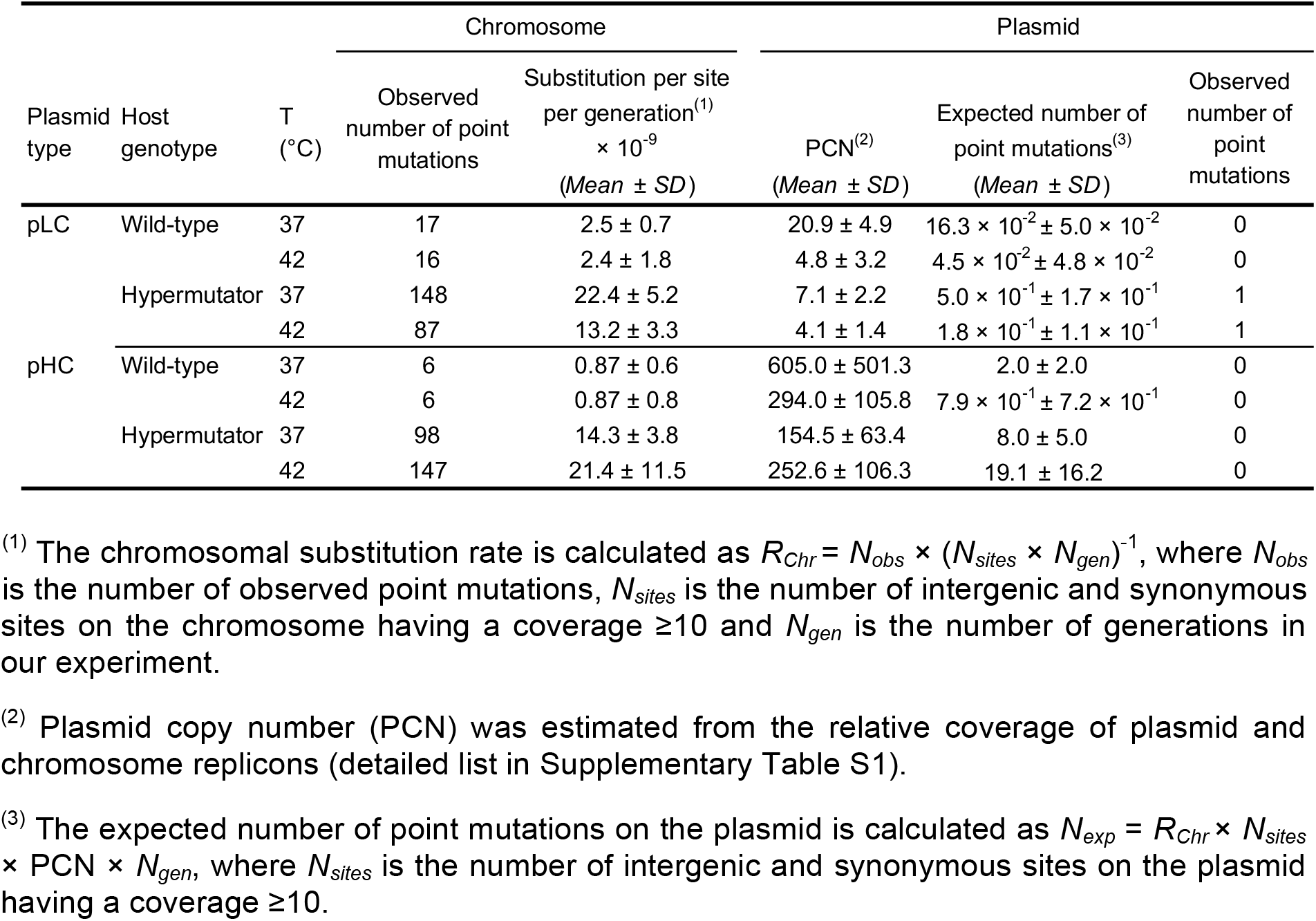
Number of intergenic and synonymous point mutations detected in the subsampled read data (104x). A detailed information used for the calculation of the mean and standard deviation for each of the eight experimental factor groups is shown in Supplementary Table S1. Additional low frequency point mutations were detected on the pLC plasmid considering the full sequencing coverage (see variant details in Supplementary Table S2). A detailed list of all observed variants on the chromosome is given in Supplementary Table S3. The variant distribution across the replicons is depicted in Supplementary Fig. S1. We note that plasmid copy number estimate from the relative sequencing coverage of small plasmids may differ from the estimated PCN using qPCR due to the DNA extraction method as well as the sequencing approach (Becker et al. 2016). The large-scale estimates of PCN from sequencing results are lower than those obtained with qPCR, hence our estimate of the plasmid mutational supply is conservative. The ancestral PCN estimated from the sequencing results are as following: pLC-wt: 2, pLC-hypermutator: 13, pHC-wt: 86, pHC-hypermutator: 116. A comparison of the PCN distribution in the evolved populations reveals that the PCN is significantly different from that of the ancestor plasmid in several of the experimental evolution settings. However, the direction of change (i.e., PCN increase or decrease) is heterogeneous (see supplementary table S1 for details).

The number of point mutations on the pHC plasmid was further validated by the analysis of the total pHC population sequencing data (i.e., samples were prepared without selection for plasmid hosts). Considering the full sequencing coverage, we detected one additional point mutation on the plasmid. The intergenic point mutation was observed with a low allele frequency of AF = 0.04 (Supplementary Table S2) in one of the pHC populations having the hypermutator host genotype. In addition, we compared the number of point mutations observed on the chromosome in the pooled pHC host populations to that of the total population sequencing data (Supplementary Table S1). The comparison shows that the wild-type populations include, on average, two additional point mutations per population in comparison to the corresponding data of pooled plasmid host colonies. The comparison of hypermutator populations shows that the total population sequencing data includes, on average, 14 point mutations per population less than the pooled colonies sequencing data. Overall, the above comparison between the two sequencing approaches demonstrates that the number of point mutations observed in the pooled host colonies data is representative both for the number of variants on the pHC plasmid and the chromosome.

This discrepancy between the observed and expected frequency of point mutations on the pHC plasmid can be explained under the segregational drift hypothesis as due to the rapid loss of most plasmid mutations. Our analysis further reveals that the allele frequency of plasmid variants is considerably lower in comparison to that of chromosomal variants. Of the total chromosomal point mutations, 30% reached fixation within the population (AF ≥ 0.9), whereas point mutations on plasmids remain at a low frequency (AF ≤ 0.09; Supplementary Fig. S2). The stark difference in the allele frequency distribution between the plasmid and chromosomal alleles indicates that the increase in plasmids allele frequency is considerably slower in comparison to chromosomal alleles. This observation is in agreement with our hypothesis according to which mutations that emerge on multicopy plasmid genomes are repeatedly ‘diluted’ in the population due to segregational drift.

### Plasmid population dynamics

What are the theoretical expectations for the impact of segregational drift on the dynamics of plasmid alleles in comparison to chromosomal alleles? To address this question we simulated plasmid and chromosome allele dynamics. For this purpose, we applied the standard haploid Wright-Fisher model to simulate forward-time evolution of a population of cells that is subject to random genetic drift (Gillespie 2010). In the simulation, the cell population size remains constant throughout evolution. To simulate plasmid evolution, we added an additional layer of random genetic drift by applying a Wright-Fisher model to the plasmid inheritance (similarly to mitochondria simulated evolution (Peng and Kimmel 2005)). Thus, during cell division, plasmid alleles are selected randomly into the daughter cells of the next generation. In terms of population genetics, the plasmid population in the simulated evolution has a substructure within the population. This implies that plasmid allele dynamics have two hierarchical levels: within the cell and within the total population.

We compared the allele frequency dynamics of neutral variants emerging in a population of *N* = 10^5^ cells on a haploid chromosome (termed C×10^5^) and two multicopy plasmids: one plasmid has a copy number of 10 plasmids per cell (termed p10×10^5^), while the second plasmid has 100 copies per cell (termed p100×10^5^). The plasmid copy number remains constant throughout the simulation, hence, a population of 10^5^ cells harbours a total of 10^6^ p10 replicons. The total number of p100 replicon in such a population is 10^7^. In order to examine the effect of the increased total number of plasmid replicons, we simulated in addition the allele dynamics of a haploid chromosome with two additional population sizes. In the first population, the number of cells equals the total p10 replicons, i.e., *N* = 10^6^ (termed C×10^6^). In the second population the number of cells matches the total p100×10^5^ replicons, i.e., *N* = 10^7^ (termed C×10^7^). In addition, we eliminate the effect of replicon size on the probability of mutation by defining the plasmids and chromosomes to have an equal size of 500 loci. The evolution of all replicons was simulated over 1,000 generations in 100 replicate populations. The simulation results reveal that the number of accumulated mutations on the plasmids is higher in comparison to C×10^5^ and lower in comparison to chromosomes with the respective population size (i.e., p10×10^5^ vs. C×10^6^ and p100×10^5^ vs. C×10^7^; Fig. 2A). These results demonstrate that the number of accumulating plasmid mutations is lower than the expected number according to plasmid mutational supply (i.e., total plasmid replicons). This discrepancy indicates that most plasmid mutations are lost rapidly after their emergence. Further comparison of the allele frequency distribution among replicons shows that the median AF of p10×10^5^ and p100×10^5^ is smaller in comparison to C×10^5^ and in comparison to the chromosome of the respective population size (Fig. 2B). This demonstrates that the increase in allele frequency of plasmid mutations is generally slower in comparison to the mutations emerging on a haploid chromosome.

**Figure 2.**
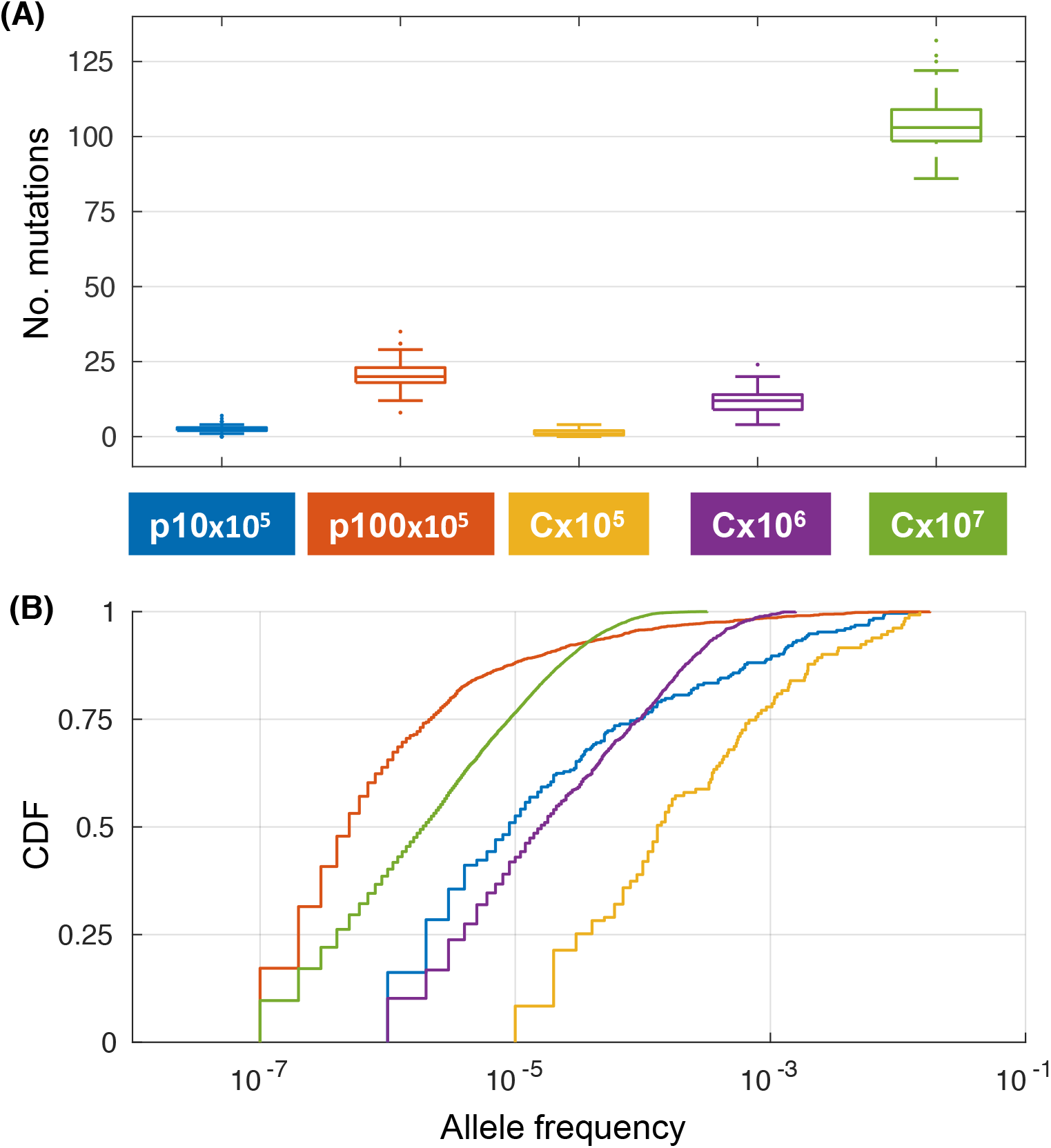
Simulated evolution of plasmids and chromosomes under neutrality with mutation (500 loci, mutation rate 2 × 10^-9^ per locus per generation). (**A**) Distribution of the number of mutations after 1,000 generations in 100 replicates. The color of the boxes corresponds to the combination of replicon and population size as in the legend. (**B**) Joint cumulative distribution function (CDF) of the allele frequency of all mutations present in the replicates after 1,000 generations.

To compare the dynamics of mutant alleles residing on the plasmid or chromosome while excluding the effect of mutational supply, we simulated the inheritance of a mutant allele already present in the population at a specified initial frequency (*f*). Thus, the allele dynamics in the population are determined by the combination of the initial mutant allele frequency, random genetic drift and the mutant allele selection coefficient (*s*). The simulation of an initially low-frequency mutant allele (*f* = 0.0001) that is neutral to the fitness of the cell (i.e., selection coefficient *s* = 0) shows that the majority of mutant alleles on plasmids are lost quickly in comparison to mutant alleles on chromosomes (Fig. 3). For example, the loss of the mutant allele in 20% of the replicates was observed at generation 22 for p10×10^5^ and at generation 76 for p100×10^5^. The chromosomal replicons lost 20% of the mutant allele much later: generation 128 in C×10^5^, generation 139 in C×10^6^ and generation 1,136 in C×10^7^ (Fig. 3A). Increasing the initial mutant allele frequency 10-fold (*f* = 0.001) results in a similar rate of allele loss between the plasmids and C×10^5^ populations. Nonetheless, loss rates of plasmid mutant alleles remain higher in comparison to chromosomal mutant alleles in simulations with the equivalent total number of replicons (Fig. 3, *s* = 0, *f* = 0.001; Supplementary Table S4). Overall, our results indicate that the loss dynamics of plasmid alleles due to segregational drift depend on the initial mutant allele frequency in the total population and the distribution of the mutant allele among plasmid hosts. Notably, the initial allele frequencies used in the simulation may be encountered upon the acquisition of a new plasmid variant in the population, e.g., by conjugation or transformation. In the case of *de novo* plasmid mutations, the initial allele frequency of the mutant plasmid is expected to be extremely low. Hence, according to our simulations, in the absence of a strong benefit for the cell, a mutation on a plasmid will be quickly lost.

**Figure 3.**
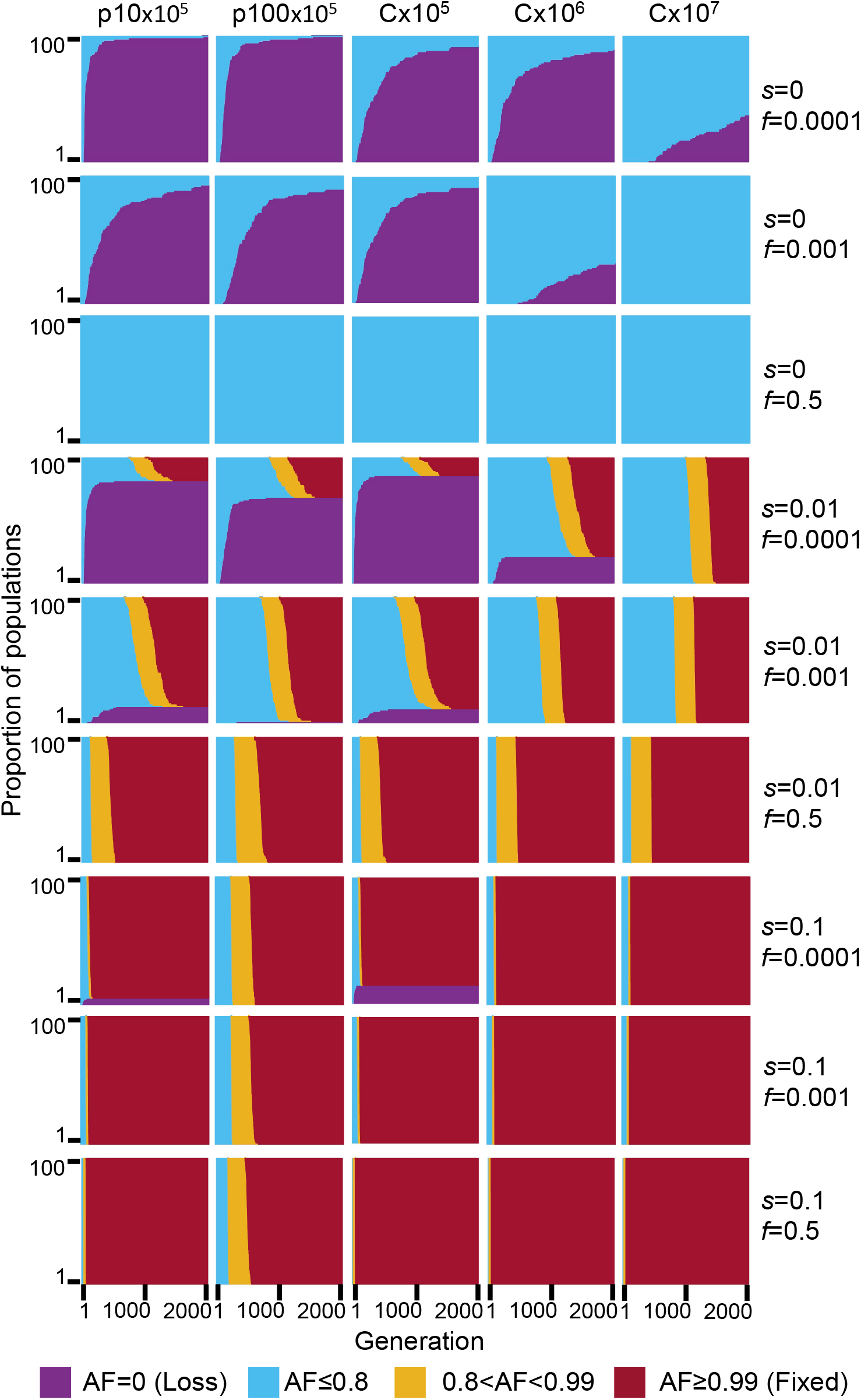
Simulated evolutionary dynamics of mutant alleles on plasmids and chromosomes. Allele frequency dynamics over 2,000 generations in 100 replicate simulated populations. (**A**) Columns are the five combinations of simulated replicon type and population size, and rows are the different combinations of selective coefficient (*s*) and initial allele frequency (*f*). Generations are on the x-axis, proportion of AF of replicates are stacked along the y-axis, population allele frequency is colour coded. For f=0.001 and f=0.0001, the starting frequency of mutant alleles is randomly sampled with the respective frequencies. For f=0.5, the starting frequency of the mutant allele is exactly 0.5 for the chromosome and exactly 0.5 inside each cell for the plasmids. Note that, the initial frequency of cells with a mutant allele in the population is higher in the plasmid simulations compared to chromosome simulations (see Supplementary Table S4 for expected number of mutant cells).

Performing the simulations where the allele has a positive selection advantage (*s* = 0.01) reveals a higher loss rate of the mutant allele in p10×10^5^ in comparison to p100×10^5^, yet the median fixation time is lower for p10×10^5^ than for p100×10^5^. Notably, the median fixation time of plasmid mutant alleles that are not lost is smaller in comparison to chromosome populations of comparable total replicon numbers when the initial frequency is low albeit the majority of plasmid populations lost the mutant allele under these conditions (Fig. 3, *s* = 0.01, *f* = 0.0001; Supplementary Table S4). The median fixation time of plasmid mutant alleles is similar to the chromosomal one when the initial frequency is increased (Fig. 3, *s* = 0.01, *f* = 0.001; Supplementary Table S4). With a high selection coefficient (*s* = 0.1), mutant allele loss is smaller than for *s* = 0.01 and the median fixation time of p10×10^5^ is similar to the chromosome populations. Notably, the median fixation time in p100×10^5^ is substantially higher in comparison to all other replicons under these conditions (Fig. 3, *s* = 0.1, *f* = 0.0001 and *f* = 0.001; Supplementary Table S4).

In the presence of balancing selection for multiple plasmid alleles (e.g., alleles supplying resistance for different antibiotics), plasmid loci may remain polymorphic over a long time-scale. Previous studies showed that when the balancing selection is replaced by selection for only one of the plasmid alleles, alternative plasmid alleles may remain polymorphic in the population for a certain time (Bedhomme et al. 2017; Rodriguez-Beltran et al. 2018). We hypothesize that the persistence of alternative plasmid alleles in the population in the absence of selection for the allele, is affected by segregational drift. To test our hypothesis, we performed simulations of plasmid allele dynamics post balancing selection. Thus, the initial mutant allele frequency is set to exactly *f* = 0.5 (i.e., each cell has equal proportions of the ancestral and mutant allele). Here we compare the dynamics of plasmid alleles to chromosomal alleles with the same total replicons as in the above simulations. Our results show that in the absence of selection for the mutant allele, both the ancestral and the mutant allele remain in the population in equilibrium as expected (Fig. 3, *s* = 0, *f* = 0.5; Fig. 4A, *s* = 0, *f* = 0.5). When the mutant allele is beneficial (*s* = 0.01 or *s* = 0.1), the ancestral allele persists in the population when it is on the plasmid for a longer time scale in comparison to a chromosomal allele of the same total replicon population (Fig. 3, s=0.01 and s=0.1, f=0.5). Nonetheless, the proportion of mutant cells in the population declines at the beginning of the simulation, hence, the beneficial mutant allele is tends to be lost in cells – also when the mutant allele is beneficial (Fig. 4A, *s* = 0.01 and *s* = 0.1, *f* = 0.5). Furthermore, the consequences of segregational drift to the loss of the mutant (beneficial) allele are more pronounced in the low copy plasmid (p10) simulations. The simulations of the high copy plasmid (p100) with *s* = 0.1 show that the ancestral allele may persist in the population four-fold longer than the expected for a chromosomal allele with the same total number of replicons (i.e., Cx10^7^; Fig. 3; Fig. 4B).

**Figure 4.**
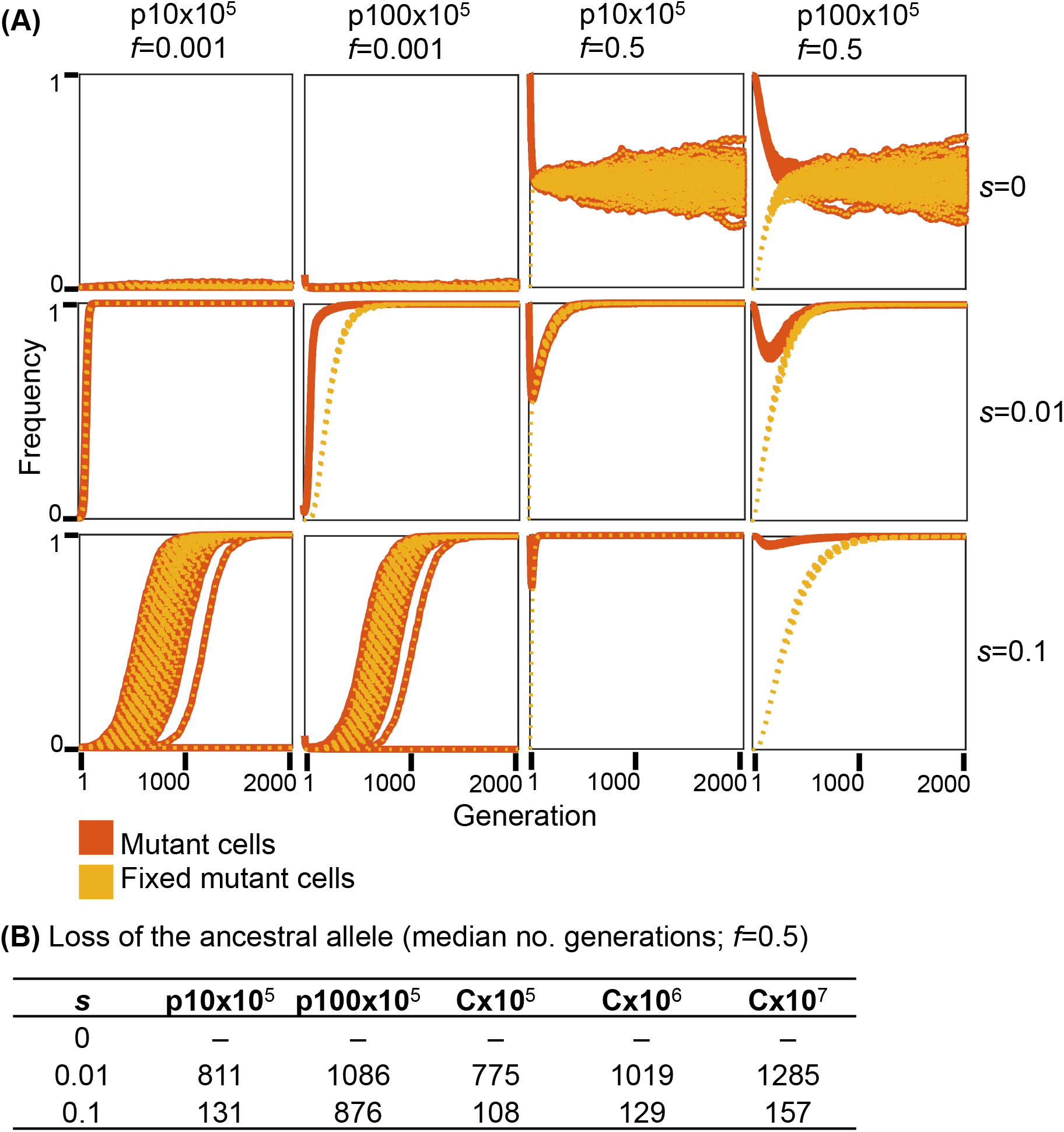
Frequency trajectories of plasmid alleles per cell in the plasmid simulations. (A) Data shown for with initial allele frequencies *f* = 0.001, and *f* = 0.5. The frequency of mutant cells is calculated as the proportion of cells with at least one mutant plasmid allele within the population. The frequency of fixed mutant cells is the proportion of cells in which all plasmids loci contain the mutant allele (i.e., the plasmid allele is fixed in the cell). Starting with a low initial allele frequency (*f* = 0.001), the difference between mutant plasmid allele dynamics and mutant cells in the population is most pronounced with *s* = 0.01. The results of the simulation where *s* = 0.1 are presented for comparison; in this setting, the dynamics of mutant plasmid alleles and mutant cells are similar. In the simulation of plasmid allele dynamics post balancing selection (*f* = 0.5) the ancestral and mutant alleles are set to an equal ratio within all cells At the beginning of the simulation. (B) Ancestral allele loss is calculated as the median number of generation where the ancestral allele was lost across all simulated populations.

The longer fixation time of p100×10^5^ alleles under the simulated conditions of strong selection (s=0.1) are well explained by the effect of segregational drift on the rate of plasmid allele fixation within the host cell. The high selection coefficient acts in favour of cells harbouring at least one plasmid mutant, such that those cells rise quickly to fixation within the population. In comparison, the increase of the plasmid allele in the total population is slower due to the effect of random genetic drift of plasmid alleles during cell division, which hinders the increase of plasmid allele frequency in daughter cells. In other words, when the plasmid copy number is high, plasmid alleles within the cell remain polymorphic (i.e., heterogeneous) over a longer time scale, also when the mutant allele is beneficial. This is also observed for the post balancing selection simulation where the loss of the non-mutant (less beneficial) alleles is prolonged for higher copy numbers.

Notably, a plasmid copy number of 100 is within the range of PCNs observed in natural isolates (Projan et al. 1987; Zhong et al. 2011), hence, the extended fixation (and loss) time observed for p100×10^5^ is likely to occur also under natural conditions. Since our simulations do not include intra-host plasmid recombination, the observed differences in allele dynamics between plasmids and chromosomes are solely due to inheritance and segregational drift. Overall, the results of the simulation demonstrate that segregational drift of multicopy plasmids has a considerable impact on plasmid allele dynamics in the population by increasing the loss rate and fixation time of plasmid alleles.

### Evolution of the host chromosome

The comparison between the evolved and ancestral populations revealed that most of the evolved genetic variants are found on the chromosome (Table 1 and Supplementary Table S3). The majority of the chromosomal variants (90%) in our experiment are observed in the hypermutator populations where the substitution rate is, on average, 10-fold higher in comparison to the wild-type populations (Table 1). Genetic variants are observed in 638 protein coding genes and ncRNA genes, of which 173 (27%) genes accumulated genetic variants in more than one population (Fig. 5A). Of the mutated genes, 60 genes (35%) are specific to populations belonging to one of the eight experimental factor combinations, indicating that the combination of host genotype, plasmid replicon type and temperature had an impact on the evolution of these genes. Furthermore, a total of 50 mutated genes (29%) are shared among hypermutator populations having the same plasmid type, regardless of the growth temperature (Fig. 5A). Hypermutator-pLC populations share, on average, 19.62 ± 2.09 (*SD*) mutated genes, whereas hypermutator-pHC populations show a lower level of parallelism with 14.45 ± 3.30 shared genes on average (SD). Furthermore, two small clusters of parallel mutated genes are observed in the wild-type hosts, the first cluster comprises the pLC populations and the second cluster includes the pHC populations evolved at 42°C.

**Figure 5.**
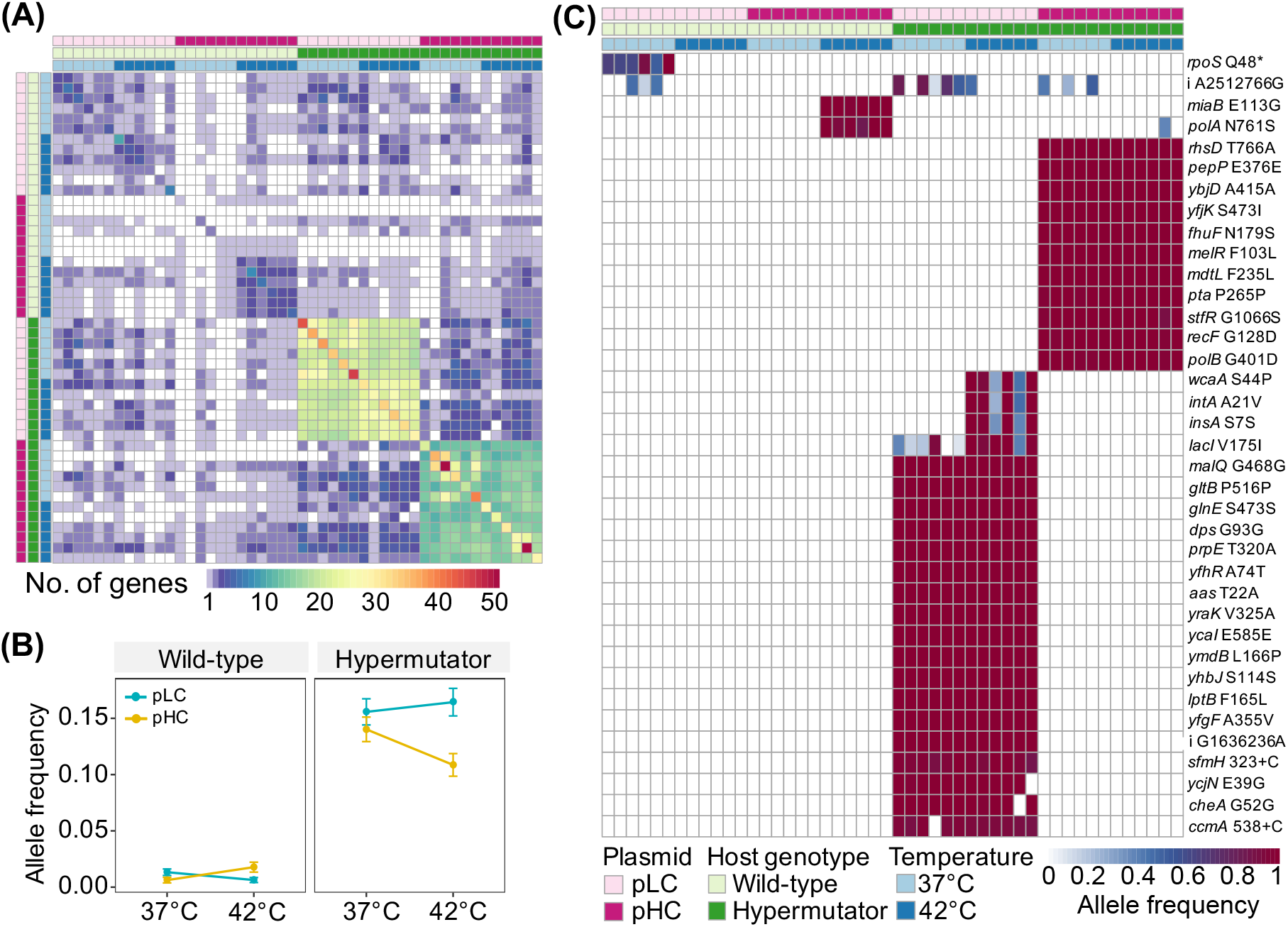
Parallel evolution of the chromosome. (**A**) A heatmap representing the number of shared mutated genes among the 48 evolved populations. The number of genes mutated in population pair *x*, *y* is calculated as the number of genes in which genetic variants (single base substitutions or indels) are observed in both populations. Cells along the diagonal present the number of genes where a mutation was detected in the corresponding population and at least one additional population. Annotation bars at the top and left right present the experimental factors (color-coded as shown in (C)). (**B**) An interaction plot of the main factors in the experiment as calculated from the complete set of allele frequencies of parallel variants (included in Supplementary Table S3). Dots represent means and error bars represent the standard error of the mean. ANOVA demonstrated significant effects for plasmid replicon type and temperature along with significant interactions between the three factors used in the experiment (statistics in Supplementary Table S5). (**C**) Color-coded matrix of variants whose allele frequency is significantly different among the main factors or their combination (using ANOVA on aligned rank transformed allele frequencies and FDR; statistics in Supplementary Table S6). Intragenic variants (11 synonymous and 22 non-synonymous) are indicated by the gene symbol and, in case of a single base substitution, coded as [ancestral amino acid][amino acid position][alternative amino acid]. The two insertion mutations are indicated by a gene symbol and coded as [nucleotide position in gene][+ inserted nucleotide]. [^*^] indicates a non-sense codon. The two intergenic variants are indicated by [i] coded as [ancestral nucleotide][genomic position][alternative nucleotide].

At the resolution of specific mutations, 144 (15%) variants were detected in more than one of the 48 evolved populations. The allele frequency of variants occurring in more than one population is significantly higher than the allele frequency of population specific variants (*P* < 0.001, using two-sample Kolmogorov-Smirnov test, *n_parallel_* = 720, *n_specific_* = 841). To quantify the contribution of the three experimental evolution factors to parallel evolution of the host chromosome, we compared the allele frequency of parallel genetic variants among all populations. We observed a significant effect of plasmid type (*P* = 1.1×10^-69^) and temperature (*P* = 3.9×10^-46^) on the allele frequency variation (using ANOVA on aligned rank transformed allele frequencies (Wobbrock et al. 2011); detailed statistics in Supplementary Table S5). In the wild-type populations, there is a strong interaction between plasmid type and temperature, whereas in the hypermutator populations, the same interaction is observed along with a main effect of the plasmid type (Fig. 5B). Our results thus reveal that the effect plasmid replicon type on the chromosomal allele dynamics is equal or larger in comparison to the growth temperature. Elevated growth temperature has an impact on *E. coli* physiology and consequences to its evolution (Deatherage et al. 2017). A recent study showed that plasmid acquisition may have a physiological effect on its *Pseudomonas aeruginosa* host (San Millan et al. 2018). Our results suggest that the effect of plasmids on the host metabolism and evolution can be equal or larger than that of the environmental temperatures.

To further identify variants that are characteristic for specific factor combinations, we tested for the effect of the factors and their interaction on the allele frequency of each parallel variant (using ANOVA on aligned rank transformed allele frequencies and FDR). This results in 37 (2.7%) variants whose allele frequency is significantly different among the factor combinations (Fig. 5C; Supplementary Table S6). A single variant occurred in all wt-pLC populations evolved at 37°C, where a nonsense mutation of Glutamine into a stop codon occurred in the *rpoS* gene. Hence, that gene is likely non-functionalized in these populations. The *rpoS* encodes the RNA polymerase σ^S^-subunit that is the master regulator of the general stress response in *E. coli* (e.g., (Patten et al. 2004)). Loss of RpoS function was previously reported to occur rapidly in nutrient-limited chemostat populations (Notley-McRobb et al. 2002). A cluster including two non-synonymous substitutions is observed for wt-pHC populations evolved at 42°C. The substitutions occurred in a gene encoding for a tRNA modification enzyme (*miaB*) and a DNA polymerase I gene (*polA*). Overall, we identified 11 different *polA* mutations in 16 pHC-populations and none in pLC-populations (Supplementary Table S3). Mutations in *polA* have been previous reported to decrease the copy number of high copy number ColE1 plasmid mutants in *E. coli* (Yang and Polisky 1993). Most of the *polA* mutations in our set are located in the same region (Klenow region), similarly to the previously reported mutants. This suggests that the *polA* mutations observed in the wt-pHC populations evolved as a response to the pHC high copy number.

Mutations that are characteristic to hypermutator-pLC populations (Fig. 5C) are found in genes that belong to four main functional categories, including carbohydrate metabolism, fatty acid metabolism, biofilm formation and stress response. Example genes where amino acid replacements have occurred are propionyl-CoA synthetase (*prpE*), a part of the propionate metabolism pathway (Textor et al. 1997), and a fimbrial-like adhesin protein (*sfmH*) that is known to promote adhesion in *E. coli* (Korea et al. 2010). Two additional non-synonymous substitutions in genes encoding for an integrase (*intA*) and a glycosyl transferase (*wcaA*) are common to all six hypermutator-pLC populations evolved at 42°C (Fig. 5C). The evolution of hypermutator-pHC populations is characterized by a majority of non-synonymous point mutations. These variants are fixed (i.e., AF > 0.9) in all hypermutator-pHC populations. The mutated genes are classified into three main categories including carbohydrate metabolism, stress response and DNA replication. Example mutated genes are DNA polymerase II (*polB*) and DNA gap repair protein (*recF*). Taken together, the differential acquisition of mutations in hypermutator populations carrying plasmid pLC or pHC suggests that the plasmid replicon type had a significant impact on the host chromosome evolution.

## Discussion

Previous experimental evolution studies of plasmid-host adaptation noted that evolved variants are mostly observed in the host chromosome rather than the plasmid genome (Bouma and Lenski 1988; San Millan et al. 2014; Loftie-Eaton et al. 2017), and this is indeed the case also for our study. Naturally, the probability for a mutation in a chromosome of the size of *E. coli* (4.6 Mb) is higher than that of our model plasmid genomes. Consequently, more mutations are likely to arise on the chromosome than on the plasmid. Notably, in most studies of plasmid-host coevolution (including ours), the selection pressure is imposed on the maintenance of the plasmid in the host population rather than a specific plasmid allele (e.g., as in (San Millan et al. 2016)). Because plasmids depend on the host machinery for their replication and persistence, the selection for plasmid maintenance is, in practice, imposed on the host machinery rather than on the plasmid itself. Moreover, the potential adaptive responses to the selective pressure for the plasmid maintenance are numerous in the complex host chromosomes (e.g., (San Millan et al. 2014; Harrison et al. 2015)), but are very limited in the genetically compact plasmid genome (e.g., (Porse et al. 2016)). This, in combination with the higher probability for mutations on the chromosome, implies that most variants - adaptive or neutral - are expected to occur on the chromosome. This can explain the high frequency of genetic variants observed on the chromosome in our experiment, as well as in other studies (e.g., (San Millan et al. 2014; Harrison et al. 2015; Loftie-Eaton et al. 2017)).

The significant effect of plasmid type on the chromosomal allele frequency observed in our experiment (Figs. 5A,C) suggests that evolution of the host chromosome depends on the replicon type of our model plasmids, with the origin of replication being a prominent characteristic thereof. The evolutionary scenario portrayed here is highly relevant for plasmid-mediated evolution of antibiotics resistant strains (e.g., (Sandegren et al. 2011; Jechalke et al. 2014; Stoesser et al. 2016)) as in such incidents, the antibiotics impose a selection pressure for the plasmid maintenance in the population. Our findings suggest that the evolution of plasmid-mediated antibiotics resistance is expected mostly to involve the host adaptation to the plasmid replicon type rather than evolution of the plasmid encoded trait.

The results from our experimental evolution of model plasmids demonstrate that the low-copy replicon has accumulated more point mutations in comparison to the high-copy replicon (Table 1). This observation contrasts with an expectation derived solely from mutation rates and genome size. This deviation from the naive expectation can be explained by the inverse correlation between population size and the impact of genetic drift on allele dynamics in the population. The low-copy plasmid constitutes a small replicon population within the cell. The dynamics of allele frequency due to drift in small populations are expected to have large fluctuations such that novel alleles are quickly lost or fixed (Bodmer and Cavalli-Sforza 1976). In contrast, the high copy plasmid constitutes a large replicon population within the cell where fluctuations in allele frequency over time are expected to be small (Bodmer and Cavalli-Sforza 1976). The impact of segregational drift on plasmid alleles is best understood in comparison to the chromosomal alleles. While novel chromosomal alleles emerging on a haploid chromosome are inherited to all daughter cells in the population, the inheritance of plasmid alleles depends on the segregation pattern of plasmid copies. Indeed, our simulations demonstrate that the frequency of neutral mutations emerging on a multicopy plasmid is higher in comparison to a haploid chromosome (Fig. 2), which is in agreement with the higher mutational supply of multicopy plasmids (San Millan et al. 2016). Nevertheless, the median allele frequency of mutations arising on multicopy plasmids is lower than that of chromosomal mutations in a comparable population size (i.e., number of replicons). Hence, the allele frequency of mutations residing on multicopy plasmids is increasing slower in comparison to chromosomal alleles. Following the emergence of a novel plasmid allele, plasmid segregation during cell division has three potential outcomes for the plasmid genotype of the daughter cells: 1) loss of the novel plasmid allele, 2) the presence of polymorphic plasmid alleles in the cell, or 3) fixation of the novel plasmid allele in the cell (see illustration in Fig. 6). The expected effect of plasmid copy number on the allele dynamics is best observed in the simulated evolution of beneficial alleles (i.e., s > 0): on the low-copy plasmid (p10), the loss of plasmid alleles occurs more frequently in comparison to the chromosome, but the median fixation time is nearly the same as that of the chromosome. Indeed, previous studies showed that under strong selective conditions, beneficial mutations on multicopy plasmids can rise quickly to fixation (San Millan et al. 2016). Notably, our simulation of beneficial alleles reveals that alleles on the high copy plasmid (p100) remain polymorphic over a longer time scale and have a higher median fixation time (Fig. 3). For neutral alleles, segregational drift of multicopy plasmids hinders the increase of plasmid allele frequency; it leads to frequent and rapid mutant allele loss on the one hand, and to longer fixation time on the other hand. Altogether, our results demonstrate that the effect of segregational drift on plasmid allele dynamics counteracts the effect of higher mutational supply accompanying increased copy number. Consequently, the evolvability of low-copy plasmids is higher than that of high copy plasmids despite the latter higher mutational supply.

**Figure 6.**
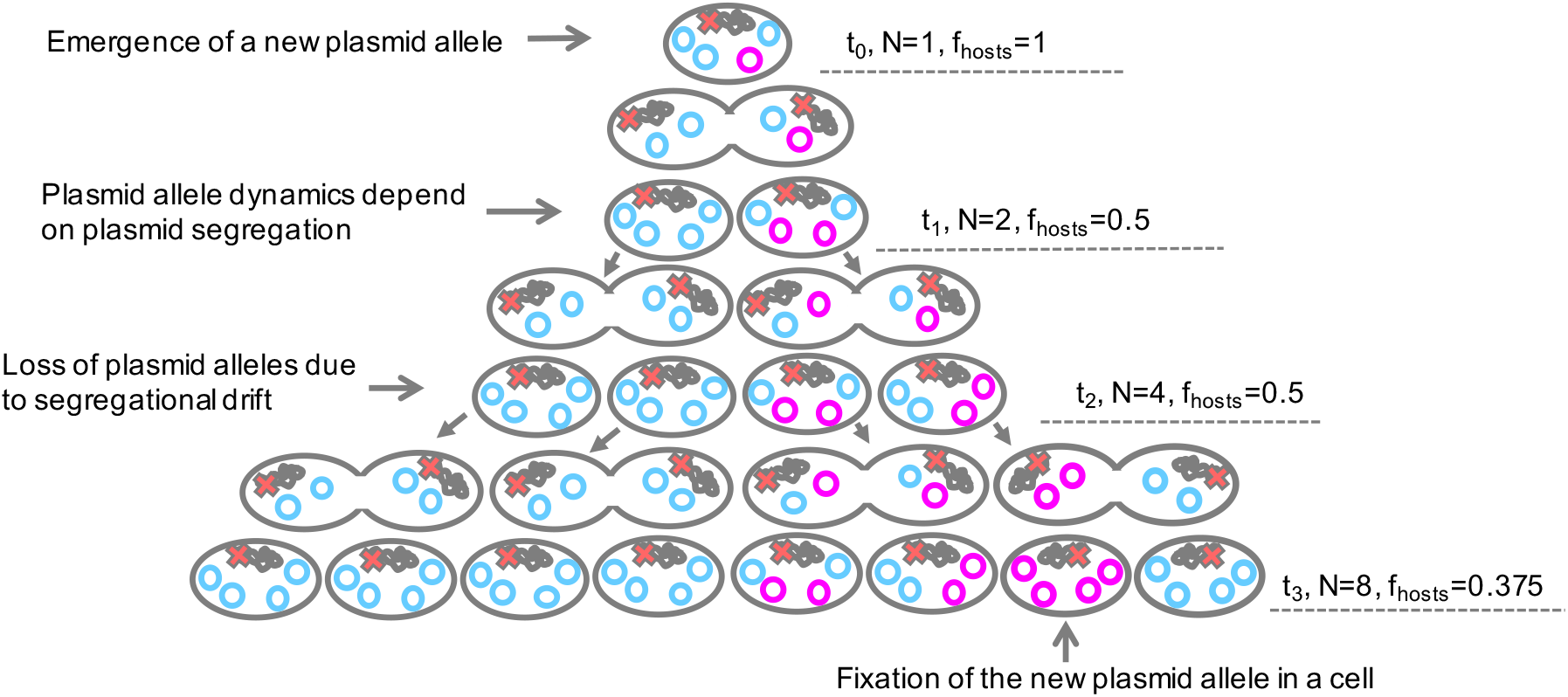
An illustration of the consequences of segregational drift to plasmid allele dynamics. The illustrated plasmid copy number is four and its segregation into daughter cells is balanced (i.e., the daughter cells inherit an equal number of plasmids). In the depicted scenario, one allele emerges on the chromosome (red X) and another allele emerges on the plasmid (magenta) at t_0_. Both alleles are considered neutral (i.e., *s* = 0). The population size (N) and the frequency of hosts (f_hosts_) over three generations are presented. The chromosomal allele is inherited into all daughter cells, such that it is present in the total population. Random segregation of the plasmid genotypes leads to decrease in the number of hosts. At t_3_, only a small minority in the population harbours the new plasmid allele. In the absence of selection for the plasmid allele presence, the new plasmid allele is deemed to remain at a very low frequency within the population (or be lost). Note: this is a simplistic example that does not include loss of host cells due to drift and it does not include consideration of recombination among plasmids.

The impact of segregational drift is not expected to be limited to non-mobile plasmids as mobile (or conjugative) plasmids are also found in multicopy state (e.g., (Figurski and Helinski 1979)). Moreover, plasmid copy number may be heterogeneous within the population, especially for plasmids that were modified towards a high copy number such as the pUC origin (e.g., (Münch et al. 2015)). Furthermore plasmid copy number can be variable during evolution (e.g., (San Millan et al. 2016; Santos-Lopez et al. 2017)). In our simulation we demonstrate that segregational drift during the evolution of plasmids having a constant copy number has an effect on their evolutionary rate. This scenario is relevant for naturally occurring plasmids having a tightly controlled copy number and segregation mechanisms. Variable plasmid copy number within the population, and over time, is expected to have implications for the effect of segregational drift on plasmid evolution and may thus lead to evolutionary rate heterogeneity in plasmid genomes. The evolution of multicopy plasmids entails time spans of intra-cellular heterogeneity, which is a situation also encountered by mitochondrial genomes within single cells. Our study thus highlights the operation of analogous evolutionary processes in bacterial plasmids and eukaryotic organelles. In summary, a realistic portrayal and modelling of plasmid genome evolution requires deeper understanding of segregational drift.

## Methods

### Bacterial strains, growth conditions and DNA techniques

*Escherichia coli* K-12 substr. MG1655 (DSM No. 18039, DSMZ) and a mismatch-repair deficient *E. coli* MG1655 derivative strain (**∆***mutS*) were used in the evolution experiment. *Escherichia coli* DH5α (Hanahan 1983) was used during construction of plasmids. All strains were routinely grown at 37°C in LB medium supplemented with 100 mg L^-1^ ampicillin when required. The evolution experiment was performed in M9 medium (M9/amp_100_) prepared as described (Sambrook and Russell 2001), supplemented with 0.4% (w/v) glucose, 0.1% (w/v) casamino acids, 20 mg L^-1^ uracil, 0.5 mg L^-1^ thiamine, 0.005% (v/v) Antifoam 204 (Sigma-Aldrich), and 100 mg L^-1^ ampicillin. Phusion polymerase (Thermo Fisher Scientific) was used for amplification of DNA-fragments for cloning and plasmid constructs were verified by restriction analysis and confirmed by sequencing. Linear DNA fragments and plasmids were introduced into *E. coli* strains by electroporation using a Bio-Rad Gene Pulser device with parameters reported previously (Dower et al. 1988). PCR-primers are listed in Supplementary Table S7.

### Construction of the host strains (wild-type and ∆mutS)

A marker-free **∆***mutS* deletion strain was constructed using the λ Red/ET Quick & Easy *E. coli* Gene Deletion Kit (GeneBridges) according to the manufacturers protocol. Briefly, the *mutS* gene was replaced with a PCR generated DNA fragment (primer pair mutS_KO_F/R) containing a kanamycin resistance marker gene flanked by FRT sites. One kanamycin resistant clone, in which the replacement of *mutS* with the marker-containing cassette was verified by PCR (primer pair PGK_F and mutS_test_R), was chosen for the removal of the kanamycin resistance marker through expression of the site specific FLP recombinase carried on plasmid 707-FLPe (GeneBridges). One resulting kanamycin-sensitive clone that had lost 707-FLPe was chosen and the deletion of the marker-containing cassette was verified by PCR using primer pair mutS_test_F/R.

In order to potentially compare mutations occurring within non-coding segments of DNA, both plasmids and host strains were equipped with segments of randomly composed, non-coding DNA. Such a 600 bp-stretch of non-coding DNA (termed art3) was commercially synthesized (GeneAart Strings Service, Thermo Fisher Scientific) and inserted into the Tn*7*-specific *attTn7* site of the wild-type and **∆***mutS* strain following the method described by (McKenzie and Craig 2006). Briefly, the art3 fragment was amplified by using primer pair art3_F/R and cloned into the *Not*I restriction site present in the mini-Tn*7*-transposon carried on the transposition vector pGRG25 (GenBank acc. no. DQ460223). The resulting plasmid was delivered into *E. coli* wild-type and the **∆***mutS* strain and a clone carrying the chromosomal *attTn7::miniTn7(art3)* insertion was verified by PCR targeting the Tn*7*-insertion site (primer pair attTn7_F/R).

### Construction of plasmids pLC and pHC

Plasmid pBBR1MCS-5 (GenBank accession no. U25061) was used to construct plasmid pLC. A 1,944-bp fragment comprising the gentamicin resistance gene and the *lac*Zα gene fragment was excised with *BstB*I and *Bsa*I. The resulting plasmid backbone fragment (2824 bp) was blunted and ligated to a blunted *Kpn*I-*Sac*I fragment containing *bla* with its promoter obtained from pBluescript SK(+) into which the *bla* fragment had been inserted into the *Eco*RV site as a blunted *Cla*I-*Not*I fragment from plasmid pKD4 (GenBank acc. no. AY048743). Next, 600 bp long stretches of random DNA (designated as art1 and art2) were inserted into the *Eco*RI site (art1) and into the *Xho*I site (art2) of pBBR1-bla, giving plasmid pLC (GenBank acc. no. MH238456). The high copy plasmid pHC (GenBank acc. no. MH238457) was constructed as plasmid pLC, but the *oriV* and the *repA* gene were then replaced by a pUC origin of replication. This was achieved by amplifying the plasmid except for the region comprising the *oriV* and the *repA* gene (using primer pair inv_pLC_F/R). The resulting fragment (2,735 bp) was treated with T4-Polynucleotide Kinase (New England Biolabs) and ligated to a PCR fragment (916 bp) comprising the pUC-origin of replication amplified from plasmid pCR4-TOPO (Thermo Fisher Scientific) with primer pair pUC-ori_F/R.

### Experimental design and setup

The evolution experiment was set up using a full factorial design of 2 plasmid types × 2 host genotypes × 2 growth temperatures. Six biological replicates were used for each of the eight experimental factor groups, resulting in a total of 48 populations that were evolved for 800 generations. The evolution experiment was conducted in chemostats made from off-the-shelf materials similar in design as previously described for the cultivation of yeast (Miller et al. 2013). The chemostat system was modified to enable bacterial culturing over longer time periods. Briefly, air breaks were introduced upstream of culturing vessels to prevent bacterial growth into medium supply lines and culture vessels were equipped with additional effluent ports to ensure constant efflux in case of clogging caused by biofilm-formation (additional information provided in Supplementary Fig. S3). Prior to the evolution experiment, plasmid pLC and pHC, respectively, were transferred into the wild-type and the ∆*mutS* strain and a single colony of each type (wt-pLC, wt-pHC, ∆*mutS*-pLC, and ∆*mutS*-pHC) was used to found a population that was serially passaged (1:1000) daily for about 100 generations in M9/amp_100_ at 37°C to acclimate the populations to the media conditions. At the onset of the evolution experiment, 50 μl overnight culture of each of the four populations (termed here ancestral populations) was used to inoculate 6 replicate chemostat cultures with a working volume of 20 ml at 37°C and 42°C each. The 48 chemostat cultures were grown to stationary densities overnight (i.e., without addition of media) and then continuously incubated at a dilution rate of 0.27 h^-1^. Hence, the growth rates are homogeneous among all populations through the growth in the chemostat. The bacterial concentration of chemostat cultures grown at steady-state was stable over time and among replicates (*M* = 1.92 × 10^8^ cells per ml, *SD* = 1.11 × 10^7^ cells per ml, *n* = 432). Nevertheless, in order to ensure the stable maintenance of flow rates during the course of the experiment, chemostat units and tubings were replaced every three weeks (approximately 200 generations). Meanwhile, chemostat populations were stored at −80°C in 25% (v/v) glycerol. From these frozen stocks, the chemostat cultures were restarted as follows: 10 ml of the frozen stock culture were pelleted by low-speed centrifugation (5000×g) for removal of glycerol and the cells were harvested in the final culturing volume of 20 ml and transferred into chemostat vessels. Each population was evolved for 90 days at steady-state resulting in approximately 800 generations. On a regular basis, all chemostat populations were reciprocally tested for cross-contamination using qPCR. To this end, each culture was tested for the presence of the *mutS* gene and for the presence of both plasmids using the following primer pairs: q_pBB_rep_F/R complementary to the *rep* gene on plasmid pLC, q_pUC_ori_F/R complementary to the *ori* on plasmid pHC, and primer q_mutS_F/R complementary to *mutS* gene in the *E. coli* MG1655 background. The proportion of plasmid bearing cells in evolved and ancestral populations was assessed by appropriately diluting samples of chemostat cultures with 1x phosphate-buffered saline, spreading on M9 agar plates and incubating at 37°C or 42°C for 14 hours. At least 100 single colonies of each chemostat sample were transferred to M9/amp_100_ and to M9 agar plates. Colonies that grew only on non-selective media were counted as plasmid-free segregants. To verify the absence of the plasmids, 1-2 randomly chosen colonies per population were chosen for testing the presence of the plasmid using qPCR (q_bla_F/R for the plasmids and q_idnT_F/R for the chromosome). The proportion of plasmid-bearing cells was determined from the comparison of the number of colonies grown on ampicillin plates to the total number of transferred colonies grown on non-selective medium.

### Plasmid copy number and loss frequency

Plasmid copy numbers were determined using quantitative PCR (qPCR) as described previously (Skulj et al. 2008), but with primers complementary to *idnT* on the *Escherichia coli* MG1655 chromosome (q_idnT_F/R) and to the *bla* gene (q_bla_F/R) on plasmid pLC and pHC. Samples of overnight cultures were heated at 98°C for 10 min and then frozen at −20°C to disrupt the cells. The qPCR reactions were carried out in a total volume of 10 μl containing 1x iTaq Universal SYBR Green Supermix (Bio-Rad Laboratories), 100 nM of each primer (final concentration) and 1 μl sample. All qPCR reactions, including positive and non-template controls, were performed in technical replicates on a CFX Connect Real-Time PCR Detection System (Bio-Rad Laboratories) using the following cycling conditions: 95°C for 3 min, and 40 cycles of 10 s at 95°C and 1 min at 59°C. Primer specificities and efficiencies were determined using standard-curve and melt-curve analyses. The ratio between the number of plasmid specific amplicons and chromosome specific amplicons is defined as the plasmid copy number (Skulj et al. 2008) and was here calculated considering the amplification efficiencies of both primer pairs.

To determine the segregational loss of plasmid pLC and pHC, five single colonies of each strain (i.e., wt-pLC, wt-pHC) were picked from M9/amp_100_ agar plates and grown overnight in 2 ml M9/amp_100_ with constant shaking at 37°C for approximately 4.4 ± 0.42 (*SD*) generations. Appropriate dilutions of stationary cell cultures were plated on M9 agar, incubated overnight at 37°C, and 100 colonies per replicate were streaked onto M9/amp_200_, M9/amp_100_, and M9 plates. The number of plasmid-free segregants was determined from the comparison of colonies grown on selective and non-selective plates.

### Whole genome sequencing

The ancestral populations of the wild-type and the Δ*mutS* strain carrying the designated plasmids (wt-pHC, wt-pLC, Δ*mutS*-pHC, and Δ*mutS*-pLC) and the 48 populations evolved for 800 generations were sequenced using Illumina sequencing technology. Population sequencing was used to enable the detection of rare variants and to asses the frequency of alleles within and across populations. Total DNA (genomic DNA plus plasmid DNA) was isolated from archived samples stored at −80°C using the Wizard Genomic DNA Purification kit (Promega). The preparation of DNA for sequencing of the evolved populations differed between the two plasmids due to the difference in their stability within the host. For pLC-populations, 1 ml of cell suspension from thawed samples was used for DNA extraction. The frequent loss of pHC led to the presence of plasmid-free segregants in the evolved populations. Therefore, in addition to the sequencing of DNA extracted from total pHC-populations, DNA was sequenced from pooled colonies of plasmid-carrying cells. The deep sequencing of plasmid-carrying hosts allows the detection of rare variants from a population of cells with a potentially heterogeneous plasmid allele pool. DNA from total pHC populations was isolated from archived samples as described for pLC-populations. The preparation of DNA from plasmid-carrying cells included a preliminary stage of selection for plasmid-hosts. Thus, frozen chemostat culture samples were first plated on LB plates supplemented with ampicillin (100 μg ml^-1^) and grown overnight at 37°C or 42°C, to select plasmid-carrying cells. Then, 100 colonies per pHC-population were pooled for extraction of total DNA (i.e., 4×600 colonies in total). Sample libraries for Illumina sequencing were prepared using either the Nextera or Nextera XT library preparation kit. All samples were quantified on a Qubit fluorometer (Invitrogen by Life Technologies) and DNA fragment length distribution was assessed on a TapeStation (Agilent Technologies). Libraries were sequenced on either a HiSeq 2500 platform with 2×125 bp reads or a NextSeq 500 platform with 2× 150 bp reads (see Supplementary Table S1 for details). Due to low initial coverage, some libraries of the wt-pLC and wt-pHC populations were re-sequenced on a HiSeq 2500 platform.

### Variant detection

Sequencing reads were trimmed to remove both Illumina specific adaptors and low quality bases using the program Trimmomatic v.0.35 (Bolger et al. 2014) with these parameters: `ILLUMINACLIP:NexteraPE-PE.fa:2:30:10 CROP:150/125` (NextSeq/HiSeq) `HEADCROP:5 LEADING:20 TRAILING:20 SLIDINGWINDOW:4:20 MINLEN:36`. Quality of sequencing reads was inspected before and after trimming using FastQC v.0.11.5 (Andrews). To confirm plasmid maps derived from partial Sanger-sequencing after plasmid construction, plasmid genomes of the four ancestral populations were assembled from trimmed paired end sequencing reads using plasmidSPAdes (SPAdes v.3.9.0 (Nurk et al. 2013)). To improve assembly speed and accuracy of the high copy plasmid pHC samples, BBNorm (BBMap tool suite v.35.82 (https://sourceforge.net/projects/bbmap/)) was used to normalize paired end read data to an average coverage of 200×. The assemblies resulted in contigs with overhangs at each end. There was no variation between plasmid assemblies of the same type. Therefore, only one assembly of plasmid pLC and pHC was used as a reference for variant detection. Plasmid genome assemblies were annotated using SnapGene software v.2.4 (GLS Biotech). The chromosomal insertion sequence art3 (described above) was included in the reference genome sequence of *E. coli* MG1655 (GenBank acc. no. NC_000913.3). Reads were then mapped to this reference genome and to the assembled plasmid genomes using BWA-MEM v.0.7.5a-r405 (Li and Durbin 2009). Corresponding mapping files of libraries that were sequenced twice were merged using PICARD tools v.2.7.1 (http://broadinstitute.github.io/picard/). Mapping statistics were retrieved using BAMStats v.1.25 (https://sourceforge.net/projects/bamstats/files/). Then, indexing, local realignment of sequencing reads, removal of read duplicates and ambiguous aligned reads were performed using PICARD tools, SAMtools v.0.1.19 (Li et al. 2009) and GATK v.3.6 (McKenna et al. 2010) retaining only paired mapped reads with a minimum mapping quality of 20. To enable a between-sample and between-replicon comparison of measures deduced from the sequencing reads (e.g., number of mutations) that is not biased by differences in sequencing coverage, subsampling of reads was performed. Therefore, mapping files were first split by contig (i.e., plasmid and chromosome) using Bamtools v.2.4.2 (Barnett et al. 2011). Reads of each chromosomal and plasmid BAM file were then subsampled to the minimal mean coverage of 104x using SAMtools (seed 1). Subsampled and non-subsampled datasets were then used to call variants. Short indels and SNPs were called using LoFreq v.2.1.2 (Wilm et al. 2012) and VarScan v.2.4.3 (Koboldt et al. 2009). Pindel version 0.2.5b9 (Ye et al. 2009) was used with options –v 30 –M 2 to detect indels. The maximum number of reads used to call variants with LoFreq and to call variants with VarScan by using SAMtools mpileup was set to 15,000,000 for non-subsampled datasets. Variants that were supported by alternative reads in one direction only were removed. Furthermore, only variants that were concordantly identified by two or more of the variant calling tools were retained using BEDTools’ intersect utility v2.17.0 (Quinlan and Hall 2010) with parameters –f 1.0 –r –u. BEDTools’ intersect utility was then used with parameters –f 1.0 –r –v to keep only variants present exclusively in evolved but not in the ancestral samples. A custom python script using pysam (https://github.com/pysam-developers/pysam) and R version 3.4.3 (R 2017) were then used to retain only variants that met the following criteria: allele frequency ≥ 0.02, depth of coverage ≥ 10, ratio of the numbers of forward and reverse reads supporting the variant ≥ 0.25, a minimum medium mapping quality of 40, a minimum medium base quality of 30 and a minimum median length of 30 nt to the end of the reads supporting the variant. Indels within an homopolymer stretch of ≥ 8 nucleotides and indels with less than 10 reads supporting the variant were excluded. Variants whose read support at the corresponding position in ancestor samples was not sufficient to detect a variant (i.e., < 10x) were excluded. Variant genomic location and the type of mutation, i.e., intergenic, synonymous, non-synonymous, or nonsense, were annotated using an in-house Perl script. Plasmid copy numbers were inferred prior to the de-duplication as the ratio of plasmid to chromosomal mean coverage. The number of intergenic and synonymous sites from the annotated reference genome sequences that were used for the mapping of sequencing reads. Intergenic positions were extracted from the reference sequences and the total number of intergenic sites was calculated as the sum of intergenic positions whose coverage was ≥ 10x. Potential synonymous sites were determined for each position within the reference sequences. The total number of synonymous sites were then calculated for each mapping file using the Nei-Gojobori method (Nei and Gojobori 1986) considering only those positions with a minimum sequencing coverage of 10x.

### Simulation

Forward-time simulations based non non-overlapping generations with a constant cell population size were performed with python3.4 and the module simuPOP v1.1.8.3 (Peng and Kimmel 2005). Two types of cells were simulated independently. Cells with chromosomes have L haploid loci (`ploidy=1`). Cells with plasmids have a predefined plasmid copy number cp that stays constant throughout the simulation. Each plasmid has L loci, resulting in a total number of L×cp loci in each cell. This was implemented based on cp customized chromosomes (`chromTypes=[sim.CUSTOMIZED]*cp`), each with L haploid loci. Alleles were simulated as binary (`alleleType=‘binary’`). Inheritance mode for the chromosome corresponds to a Wright-Fisher model (`operator RandomSelection`()). Inheritance mode for the plasmid corresponds to a Wright Fisher model with mitochondrial inheritance of organelles (`operator RandomSelection(ops=[sim.MitochondrialGenoTransmitter`()])).

For the simulations with mutation, initial populations (t=0) were homogenous, i.e., all loci had state 0. The mutation rate in the simulation was constant with 2 × 10^-9^ per locus per generation (operator `SNPmutator(u=2e-9)`). For the simulated evolutionary dynamics of mutant alleles, we simulated the evolution of one binary locus (L=1) having a mutant allele already present in the initial population at a specified initial frequency (*f*) and no new mutation is applied during simulated evolution. Initial populations were randomly generated with a particular mutant frequency f (parameter `freq=[1-f,f]` for the method `initGenotype`). Note that this assigns the mutant allele randomly to each locus, independent of the state of the other loci in the same cell for plasmid simulations. For the simulated evolution with selection, the probability to be inherited to the next generation is proportional to the fitness value of the cell. For cells with a chromosome, a fitness value of 1+s was assigned to allele state 1 and fitness value 1 was assigned to allele state 0. For cells with plasmids, a fitness value of 1+s was assigned if at least one plasmid has allele state 1, otherwise the fitness is 1 (i.e., dominant selection mode). Selection was applied using a fitness field (`infoFields=“fitness”`) and the operator `PySelector`. Simulations were run for a particular number of generations using the `evolve` method.

### Data availability

Sequence reads of evolved and ancestral populations are found in SRA (SRA acc. SRP141152). Plasmid sequences are found in GenBank (pLC: MH238456, pHC: MH238457).

## Acknowledgments

We thank Sarah Kovarik, Mathias Krüger and Rabea Plohr for their assistance in the experimental work. We thank Maxime Godfroid and Devani Romero Picazo for critical comments on the manuscript. We thank Wolfgang Streit for providing the plasmid pBBR1. The study was supported by the European Research Council (Grant No. 281357 awarded to TD).

## Author contributions

J.I, I.M., and T.D. designed the experiment. J.I. established and performed the experimental evolution study and the comparative genomic analysis. A.K. performed the simulated evolution. P.R. sequenced the genomes. T.W. and N.H. took part in the experimental work. C.W., G.L, A.K., and T.D. took part in the computational analysis. All authors interpreted the results and drafted the manuscript.

